# Long non-coding RNA RGMB-AS1 as a novel modulator of Bone Morphogenetic Protein Receptor 2 signaling in pulmonary arterial hypertension

**DOI:** 10.1101/2022.08.27.505495

**Authors:** Md Khadem Ali, Yu Liu, Lan Zhao, Joseph C. Wu, Vinicio de Jesus Perez, Christopher J. Rhodes, Aiqin Cao, Martin R. Wilkins, Mark R. Nicolls, Edda F. Spiekerkoetter

## Abstract

This study shows that the long non-coding RNA RGMB-AS1 is upregulated in the blood and pulmonary vascular cells of PAH patient and that it regulates BMPR2 signaling. Inhibiting RGMB-AS1 increases BMPR2 signaling and improves pulmonary vascular cell function.

## To the editor

Pulmonary arterial hypertension (PAH) is a complex disease of the pulmonary circulation that currently has no cure. In many forms of PAH, impaired Bone Morphogenic Protein Receptor 2 (BMPR2) signaling has been considered as one of the key pathomechanisms - either through loss of function mutations in familial PAH or downregulation of BMPR2 expression. Yet how BMPR2 expression and signaling is regulated remains largely unclear, especially in non-genetic forms. Previously, we and others have shown that targeting BMPR2 signaling and improving the BMPR2/TGF-β disbalance might be an effective therapeutic option for PAH [1–3]. Increasing BMPR2 signaling and expression using the repurposed drugs FK506 [4, 5] and Enzastaurin [6] has proven to be a successful approach in improving experimental PAH. As these drugs do not address the underlying cause for reduced BMPR2 expression and signaling, we set out to identify BMPR2 signaling modifiers that could be targeted as an alternative way to improve signaling in PAH.

Long non-coding RNAs (lncRNAs), endogenous RNAs that are longer than 200 nucleotides in size and have no protein-coding capacity, have been shown to play a significant role in pulmonary vascular remodeling, RV remodeling and PAH [7]. Yet the role of lncRNAs in modulating BMPR2 expression and signaling and their contribution to the development and progression of PAH is unknown. Our goal was to explore the role of lncRNAs in modulating BMPR2 signaling in PAH. Through a series of clinical, experimental, and bioinformatic analyses, we identified lncRNA repulsive guidance molecule B antisense RNA 1 (RGMB-AS1) as a critical emerging player in PAH.

To identify novel lncRNAs associated with BMPR2 signaling, we performed RNAseq of pulmonary arterial endothelial cells (PAECs) treated with either the BMPR2 ligand, BMP9, or BMPR2 siRNA [8]. The positive hits were defined as those lncRNAs that responded in an opposite direction when activated with BMP9 or treated with BMPR2 siRNA. The top BMP-responsive lncRNAs were individually knocked down with siRNA and their effect was assessed on the expression of ID1, a major downstream target of BMPR2 signaling. This approach identified RGMB-AS1 as a potential BMPR2 signaling inhibitor (**Figures 1A-C**), although the RGMB-AS1 knockdown efficiency using siRNA was only about 55% (data not shown). As further confirmation, we knocked down RGMB-AS1 with either siRNA or more effectively with locked nucleic acid (LNA) gapmer in the presence and absence of ligand stimulation and assessed ID1 expression in PAECs and pulmonary arterial smooth muscle cells (PASMCs). We found that RGMB-AS1 knockdown further increased BMP9- and BMP4-induced ID1 expression in PAECs and PASMCs, respectively (**Figures 1D-F**). To determine whether RGMB-AS1 regulates BMPR2 signaling through its neighboring protein coding gene RGMB, we measured RGMB expression by qRT-PCR in PAECs and PASMCs following knockdown of RGMB-AS1 with siRNA or LNA. We found that RGMB-AS1 knockdown increased RGMB expression in PAECs and PASMCs (**Figure 1G**), suggesting that RGMB-AS1 may regulate BMP signaling partly through RGMB, a BMP co-receptor that facilitates ligand binding. As hypoxia is a crucial contributor to some forms of human pulmonary hypertension (PH) and experimental PH and is associated with decreased BMPR2 signaling, we measured the expression of RGMB-AS1, BMPR2, ID1, and VEGFA (a hypoxia-responsive gene) in hypoxic PAECs by RT-qPCR. We observed that hypoxia induced expression of RGMB-AS1 and decreased BMPR2 and ID1 expression (**Figure 1H**). To determine whether RGMB-AS1 expression was increased in PAECs from PAH patients, we re-analyzed a publicly available RNAseq data set comprising 9 healthy and 9 PAH PAECs (GSE126262, [9]). Yet we did not find significant changes in RGMB-AS1 expression between the control and PAH PAECs (**Figure 1I**). In contrast, RGMB-AS1 was increased 7-8-fold in PASMCs isolated from PAH patients (n=3) compared to healthy controls (n=3) analyzed by RNAseq (**Figure 1J,** [10]). Furthermore, RNAseq analysis of whole blood of patients with idiopathic, heritable, and drug-induced PAH (n=359) compared to age- and sex-matched healthy controls (n=72) showed a significant increase in RGMB-AS1 expression (**Figure 1K**, [11]). RGMB-AS1 knockdown with siRNA or LNA increased PAEC proliferation and decreased PASMC proliferation, as assessed by MTT assay (**Figures 1L-N**), and induced PASMCs apoptosis as assessed by caspase 3/7 activity assay (**Figure 1O**).

**Figure 1 legent.**
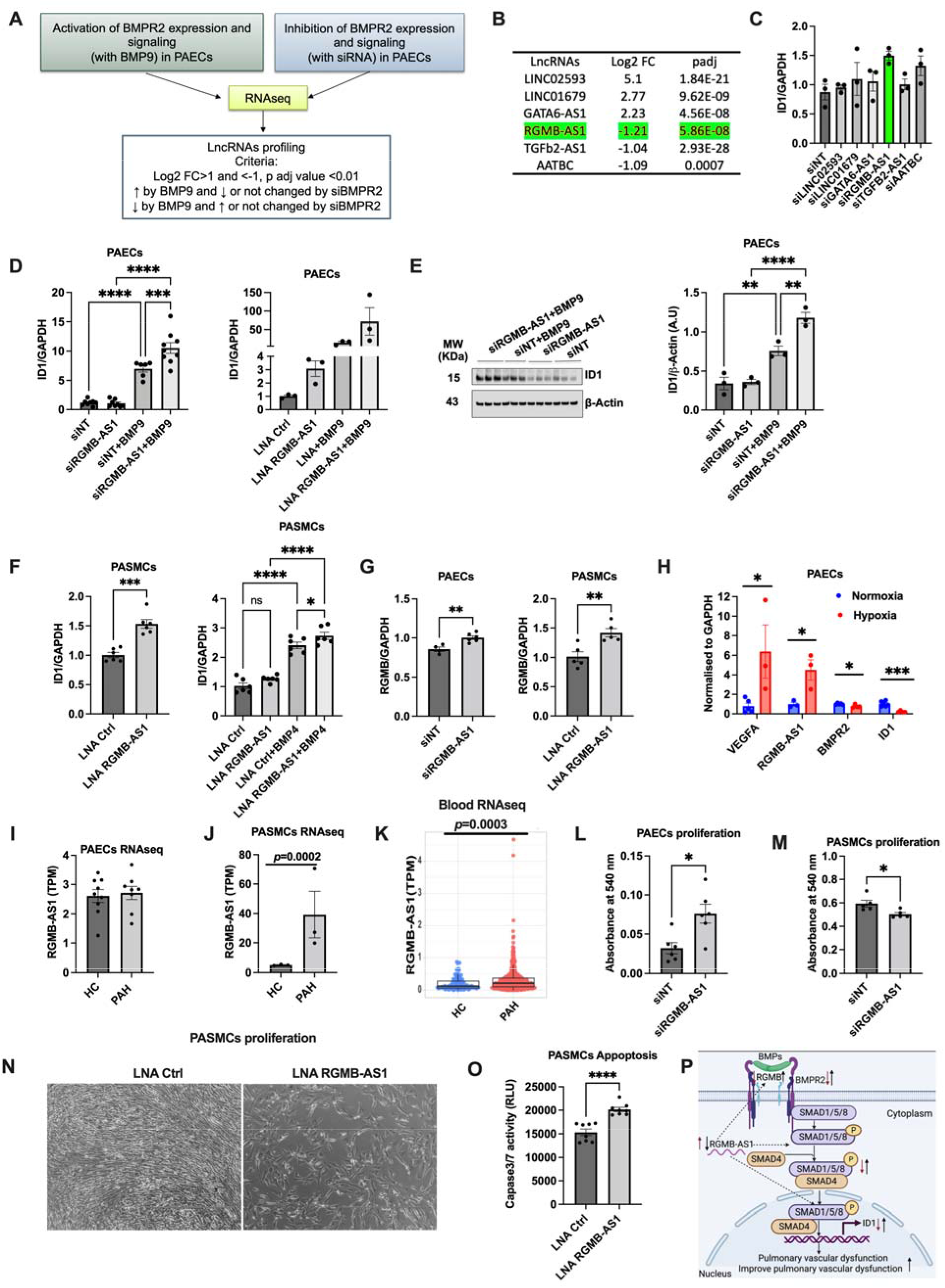
A-B) Experimental Strategy to identify lncRNAs associated with BMPR2 signaling. A) BMPR2 signaling were activated in healthy PAECs stimulated with BMP9 20 ng/ml for 2hrs and inhibited with BMPR2 siRNA transfection for 48hrs, harvested RNA and performed RNAseq. LncRNAs were screened based on the criteria that they express in an opposite direction of BMP signaling activation and inhibition condition. B) Top 3 BMPR2 activator, and inhibitor-associated lncRNAs are represented as log2 fold change from controls in the table. C-E) RGMB-AS1 knockdown increases BMP9-induced BMP signaling in PAECs. Top selected 6 lncRNAs were knocked down with siRNA and assessed ID1 expression by qPCR to see whether these lncRNAs are the upstream of the BMPR2 signaling in PAECs (C). PAECs were seeded onto 6-well plate and next day, 100nM siRNA was transfected with 2ul of RNAimax in optimum media. After 24hours, the transfected cells were starved with starvation media for 16 hours and treated with BMP9 (20ng/ml) (a ligand for BMPR2 signaling in PAECs cells) for additional 2 hours and then harvested and isolated RNA and proteins. RGMB-AS1 were also knocked down with LNA gapmers in PAECs and PASMCs. Expression of the key BMP pathway molecule, ID1 was assessed by RT-qPCR and western blotting (D-F). Densiometric analysis was performed with ImageJ. G) RGMB mRNA were measured by qRT-PCR following knockdown of RGMB-AS1 with siRNA or LNA in PAECs and PASMCs. H) RGMB-AS1 is induced by hypoxia in PAECs at 72hrs of hypoxia exposure while BMPR2 and ID1 expression is downregulated, as measured by qRT-PCR. RGMB-AS1 expression of PAECs (n=9/group) (I), PASMCs (n=3/group) (J) and whole-blood of PAH patients (n=72 healthy and n=359 PAH) (K), assessed by RNAseq. L-N) RGMB-AS1 knockdown with siRNA or LNA gapmers increases proliferation of PAECs and decreases PASMCs proliferation. PAECs and PASMCs were transfected 10nM siRNA or LNA against RGMB-AS1 with 2 ul of RNAiMax for 48 hours and assess cell proliferation by MTT assay. O) RGMB-AS1 knockdown with LNA gapmers induced apoptosis as measured by commercially available caspase 3/7 activity assay kit. P) Proposed schematic mechanism of RGMB-AS1 and BMPR2 signaling in PAH. Data are represented as mean+/-standard error mean, student t test for comparing between two groups. One-way ANOVA, LSD fisher post test test, for comparing multiple groups. *p<0.05, **p<0.01, ***p<0.001, ****p<0.001. siNT; non-target siRNA, siRGMB-AS1; RGMB-AS1 siRNA. LNA; locked nucleic acid.

Collectively, our findings suggested that RGMB-AS1 might be an important BMPR2 signaling modulator in vascular cells in PAH. However, it remains unknown how RGMB-AS1 regulates BMPR2 signaling in PH. A recent lung cancer study showed that RGMB-AS1 expression is negatively associated with the expression of its neighboring protein-coding gene RGMB [12]. Notably, RGMB is a BMP co-receptor that promotes BMP signaling in neuronal regeneration and kidney disease [13–15]. NCBI nucleotide blast search of RGMB-AS1 sequences revealed a 322nt-homologus sequence to the whole exon 2 of RGMB but in a complementary orientation (Data not shown). Our finding of a significant albeit small increase in RGMB mRNA when RGMB-AS1 was silenced in PAECs suggests that the opposite might be true as well; that increased levels of RGMB-AS1 would decrease RGMB levels, thereby reducing BMPR2 signaling (**Figure 1O**). RGMB-AS1 could also regulate BMPR2 signaling through other distant protein-coding genes, microRNAs, transcription factors, and non-canonical smad-independent pathways, such as MAPK, c-Jun, p38, AKT.

Since BMPR2 receptors are highly expressed in PAECs, we used those cells in our initial experiments to identify and validate lncRNAs involved in BMPR2 signaling. We also found a significant upregulation of RGMB-AS1 and its correlation with downregulation of BMPR2 signaling in hypoxic PAECs, in line with our findings of RGMB-AS1 as a BMPR2 signaling inhibitor. Yet, by re-analysing publicly available RNAseq data sets, we observed a significant upregulation of RGMB-AS1 in PASMCs but not in PAECs of PAH patients. Based on our RNAseq expression re-analysis studies, it seems that expression levels of RGMB-AS1 are relatively higher in PASMCs than PAECs. Because of the very low baseline expression levels of RGMB-AS1 in PAECs, there is a lot more variation in expression of the lncRNA in PAECs, which warrants further studies to confirm these findings in a larger cohort of PAH patients.

Hyperproliferation of PASMCs and abnormal proliferation and tube formation of PAECs are thought to contribute to the development and progression of PH [7]. Our *in vitro* functional assay of RGMB-AS1 knockdown data showed inhibition of PASMC proliferation, suggesting that RGMB-AS1 knockdown might be benefical in PAH by inhibiting pathological PASMC hyperproliferation. These findings are consistent with our clinical RNA expression and BMPR2 signaling data, where we observed that RGMB-AS1 is upregulated in PAH and the observation that RGMB-AS1 silencing increases BMPR2 signaling. Taken together, RGMB-AS1 modulation could influence pulmonary vascular remodeling.

While the major strength of our study is the use of an unbiased RNAseq high throughput approach, a siRNA mini-library combined with experimental validation studies to discover BMPR2 signaling associated lncRNAs that might be of clinical relevance as they are expressed in a large cohort of PAH patients, our study has some limitations. The RGMB-AS1 RNAseq expression data of PAH patients were not further experimentally validated in a second large cohort of PAH patients. In addition, we only provided RGMB-AS1 inhibition findings; it is critical to confirm whether overexpression of the lncRNA decrease BMPR2 signaling and alters cellular phenotypes, such as proliferation, apoptosis, and migration.

In summary, our results demonstrate the first piece of evidence that RGMB-AS1 is upregulated in the blood and pulmonary vascular cells of PAH patients. Furthermore, inhibiting RGMB-AS1 effects BMPR2 signaling and vascular cell function, which might be of therapeutic value to improve vascular remodeling in PAH.

## Acknowledgment

The authors thank Professor Dean Felsher (Director, Translational Research and Applied Medicine program, Stanford University, CA, USA) and Dr. Joanna E. Liliental (Associate Director, Translational Research and Applied Medicine program, Stanford University, CA, USA) for their critical comments and constructive suggestions to improve this research project. The authors thank Dr. Adam Andruska for sharing the RNAseq data set generated from PAECs treated with either BMP9 ligand or BMPR2 siRNA. The authors also gratefully acknowledge the participation of patients recruited to the the UK PAH Cohort Study consortium.

## Author contribution

MKA and ES conceptualised the study design. MKA performed the experiments and data analysis. All authors contributed to data collection, data interpretation, writing, and editing the manuscript. ES: fund acquisition and supervision.

## Abbreviations

LncRNA: long non-coding RNA
BMPR2: bone morphogenic protein receptor 2
PH: pulmonary hypertension
PAH: pulmonary arterial hypertension
PAECs: pulmonary arterial endothelial cells
PASMCs: pulmonary arterial smooth muscle cells
RV: right ventricular
RGMB-AS1: repulsive guidance molecule B antisense RNA 1
LINC02593: long intergenic non-coding RNA 2593

